# A viscoelastic-plastic deformation model of hemisphere-like tip growth in Arabidopsis zygotes

**DOI:** 10.1101/2024.07.17.603843

**Authors:** Zichen Kang, Tomonobu Nonoyama, Yukitaka Ishimoto, Hikari Matsumoto, Sakumi Nakagawa, Minako Ueda, Satoru Tsugawa

## Abstract

Plant zygote cells exhibit tip growth, producing a hemisphere-like tip. To understand how this hemisphere-like tip shape is formed, we revisited a viscoelastic-plastic deformation model that enabled us to simultaneously evaluate the shape, stress, and strain of Arabidopsis (*Arabidopsis thaliana*) zygote cells undergoing tip growth. Altering the spatial distribution of cell wall extensibility revealed that cosine-type distribution and growth in a normal direction to the surface creates a stable hemisphere-like tip shape. Assuming these as constraints for cell elongation, we determined the best-fitting parameters for turgor pressure and wall extensibility to computationally reconstruct an elongating zygote that retained its hemisphere-like shape using only cell contour data, leading to formulation of non-dimensional growth parameters. Our computational results demonstrate the different morphologies in elongating zygotes through effective non-dimensional parameters.

## Introduction

Growth patterns in cells are divided into two types: diffuse growth, where the entire cell surface grows; and tip growth, where only the tip region grows (Kropf *et al*., 1998). Characteristic features of tip-growing cells include unique cell wall properties, cytoskeleton organization, and organelle activities (Rounds & Bezanilla, 2013). Since plant cell growth is thought to be controlled by cell wall biosynthesis and orientation (Green, 1962; Redbetter & Porter 1963), quantifying cell growth patterns is important for understanding the cell wall properties behind growth, as described more precisely below. To avoid ambiguity, we define surface growth or surface elongation as extension on the cell surface and define surface point velocity as the time derivative of the displacement of the surface point. Early studies of tip growth in *Phycomyces* fungi quantified surface point velocity with tips estimated to grow at 1.2–1.4 mm per hour (Castle, 1942; Castle, 1958), while *Nitella* rhizoids were found to have a linear growth velocity of 1.7 µm/min (estimated as 0.1 mm/h) at the tip dome (Chen, 1973). During tip growth of root hairs in *Medicago truncatula* and pollen tubes in *Lilium longiflorum*, the maximal elongation zone is not located exactly at the tip but is instead located in the region slightly proximal from the tip (Shaw *et al*., 2000; Dumais *et al*., 2004; Geitmann & Dumais, 2009). These quantitative analyses indicate the importance of detailed quantification when studying tip growth in cells.

In addition to elucidating where the elongation zone is located, quantification of the direction of surface point velocity is also important. For example, imaging data from fungal hyphae with carbon particles on the cell surface revealed that some of the displacements were in directions almost perpendicular to the surface (Bartnicki-Garcia *et al*., 2000). That study inferred two important aspects using a mathematical model: (1) The driving force should be turgor pressure, and (2) the vesicle supply center should be consistent with the direction almost normal to the cell surface. Viscoelastic-plastic deformation models based on this pioneering mathematical study have been extensively applied to study tip growth (Goriely & Tabor, 2003; Dumais *et al*., 2006; Dumais, 2021). In these models, cell morphology, the mechanics on the cell surface, and the deformation of the apical region are simultaneously analyzed using mathematical equations. This allows prediction of cell shape based on the history of the mechanics and deformation of the cell. These mathematical studies have prompted other studies using mechanical models (finite element methods) with modified material properties in pollen tubes (Fayant *et al*., 2010) and the introduction of non-dimensional parameters that characterize cell shape independently of cell size for studying tip growth (Campas *et al*., 2012). Therefore, the mathematical relationships among morphology, mechanics, and deformation can be used to determine the cell wall properties during cell growth.

As described above, tip growth occurs in *Nitella* rhizoids, fungal hyphae, root hairs, and pollen tubes. We recently found that plant zygote cells also exhibit tip growth (Kang *et al*., 2023). In our previous study, we combined live imaging with the so-called normalized coordinate to show that only markers (a pollen grain expressing a sperm-specific plasma membrane) near the tip moved while other markers outside the tip region did not move. Furthermore, the elongating tip of a single zygote cell shows a characteristic hemisphere-like shape. In Arabidopsis (*Arabidopsis thaliana*) zygotes, microtubules (MTs) form a ring-like structure in the subapical region, which might support the hemisphere-like tip shape (Kimata *et al*., 2016). The hemisphere-like shape and MT ring are not typical in tip-growing cells of angiosperms, but similar structures are found in the tip-growing protonema of the southern maidenhair fern (*Adiantum capillus-veneris*) (Murata and Wada 1989). In this fern protonema, cellulose microfibrils are aligned in parallel to MTs and cell division occurs. Therefore, the zygote might utilize fern-like tip growth to produce a spherical daughter cell at the tip, which develops into a globular embryo (Ueda *et al*., 2011). Since such an approximately hemispherical shape during elongation has also been observed in the cells in fission yeast (*Schizosaccharomyces pombe*; Abenza *et al*., 2015), the existence of a unifying mechanism for shaping a spherical tip needs to be investigated.

In this study, we reimplemented the viscoelastic-plastic deformation model described by Dumais *et al*. (2006) to elucidate the mechanism that shapes the hemisphere-like tip. First, we obtained growth parameters in the model that matched the hemisphere-like shape described by the data. We determined that this shape results from surface point velocity almost normal to the cell surface around the tip region. Finally, we reconstructed the elongating zygote computationally using model parameters associated with turgor pressure and cell wall expansion derived from actual zygote cell contour data. Our findings shed light on the morphology and mechanics of tip growth in plant cells.

## Methods

### Viscoelastic-plastic deformation model

A viscoelastic-plastic deformation model was employed (Dumais *et al*., 2004; Dumais *et al*., 2006). This model comprises three steps (Fig. 1). Step 1: The stress states in the meridional and circumferential directions (*σ*_s_ and *σ*_θ_, respectively) are determined by the mechanical equilibrium of the cell with turgor pressure *P*. Step 2: The strain rates in the meridional and circumferential directions (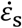 and 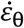, respectively) are determined by the stresses applied on the wall and the mechanical properties of the cell wall (*v*, Φ, and *σ*_y_), where *v* is the flow coupling (Dumais *et al*., 2006), Φ is the cell wall extensibility, and *σ*_y_ is yield stress. Step 3: The next shape is formulated using the previous cell shape and the velocity vectors (*v*_t_ and *v*_n_), where *v*_n_ is the velocity in the direction perpendicular to the surface, and *v*_t_ is the velocity in the meridional direction, which is one of the tangential directions.

**Fig. 1.**
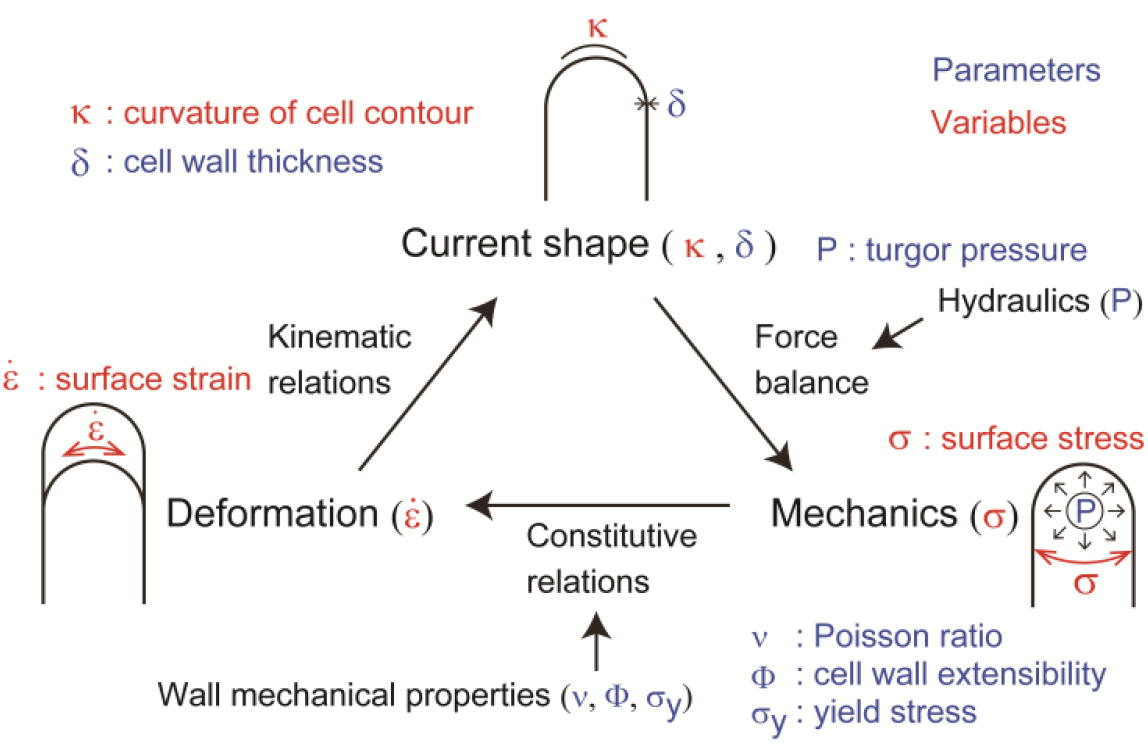
Schematic illustration of the viscoelastic-plastic deformation model. The model output is the cell contour of a tip-growing cell. By applying the hydraulics parameter (turgor pressure, *P*) to the current shape with curvature *κ* and wall thickness *δ*, the mechanics with stress *σ* are determined. By modifying the mechanical parameters of the cell wall (*v*, Φ, and *σ*_y_), the deformation with strain rate 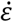 is determined.

For Step 1, the following equations were reformulated to evaluate the stress state:

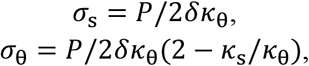

where the parameter *δ* stands for cell wall thickness, and *κ*_s_and *κ*_θ_are the curvatures in the meridional and circumferential directions, respectively.

For Step 2, the following relationship was exploited:

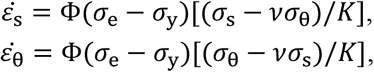

for *σ*_e_ ≥ *σ*_y_ and 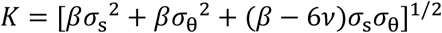, where 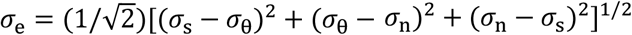 (Hill, 1998) and *β* = 2*v*^2^ − 2*v* + 2, where *β* represents the anisotropy between the two directional stresses *σ*_s_ and *σ*_θ_. Using *v*_n_ and *v*_t_, the kinematic relations for cell shape can be evaluated as follows:

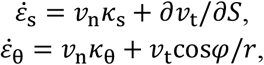

based on the cross-sectional radius *r*(*S*) with curvilinear coordinate *S*. The angle *φ* is defined as the angle between the normal vector to the surface and the axis of the cell. The normal strain rate 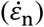 was assumed to always be zero, where thickening due to cell wall deposition (Houwink & Roelofsen, 1954; Kataoka, 1984) can be expressed in the following manner:

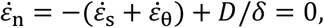

where *D* is the rate of wall deposition per unit surface area.

For Step 3, the following velocities in the normal (perpendicular) and meridional directions were obtained from the equations above:

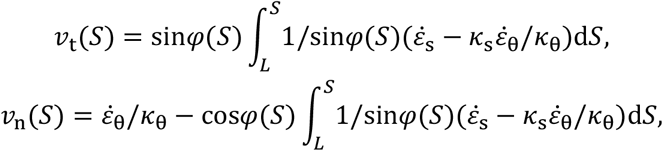

where *L* is the meridional distance between the pole and the equator. The spatial scale is based on the arbitrary unit (a.u.), which was rescaled with the typical radius scale, e.g., 5 µm.

### A hemispherical tip shape is sufficient for cosine-type wall extensibility

According to Green & King (1966) and under the model assumptions (Dumais *et al*., 2004; Dumais *et al*., 2006), a hemispherical cell shape is sufficient but not necessary for cosine-type wall extensibility, as described below. Considering a hemispherical cell shape with a constant radius *R*, the allometric coefficient (anisotropy rate), defined as the rate of the strain rate in the meridional direction to that in circumferential direction, can be expressed as

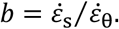

It was shown that the meridional velocity in the curvilinear direction is proportional to the *b*th power of the sine function of the curvilinear distance *S* as follows:

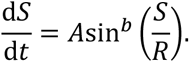

If the surface growth around the tip is approximately isotropic on the surface, characterized by *b* = 1, the tip shape remains hemispherical (Green, 1969). In the derivation, assuming that the deformation of the surface is infinitesimal, the strain rate in a small meridional element Δ*S* can be expressed as

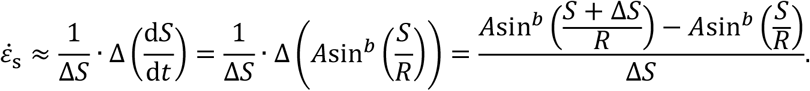

Let Δ*S* → 0, then

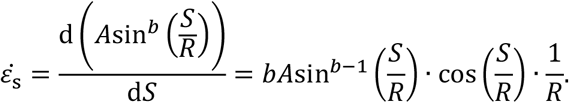

when the tip shape remains hemispherical with *b* = 1,

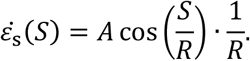

In addition, according to Dumais *et al*. (2004), the wall extensibility is estimated using the ratio

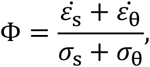

where 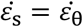 (transversely isotropic property) and *σ*_s_ = *σ*_θ_ for a hemispherical tip. To substitute the equations for *σ*_s_ and 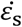 into the above expression, the following relation can be derived:

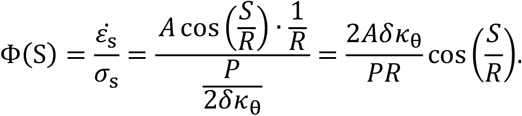

This means that a hemispherical tip geometry is a sufficient condition for cosine-type wall extensibility. Note that cosine-type wall extensibility is not sufficient for a hemispherical shape when the transversely isotropic property is not held.

## Results

### The viscoelastic-plastic deformation model can simultaneously evaluate cell shape, cell wall stresses, and strain rates

The viscoelastic-plastic deformation model enabled us to investigate wall stresses and deformation using only the cell shape (Fig. 1, Fig. 2A, see details in Methods). Specifically, we used shape information (the meridional curvature *κ*_s_, the circumferential curvature *κ*_θ_, and cell wall thickness *δ*) and turgor pressure *P* to calculate the mechanical information (the meridional stress *σ*_s_ and the circumferential stress *σ*_θ_) (Step 1). We then calculated the velocity vectors (*v*_n_ and *v*_t_) in the normal (perpendicular) and meridional directions associated with strain information (the meridional strain rate 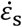 and the circumferential strain rate 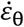) (Steps 2 and 3).

**Fig. 2.**
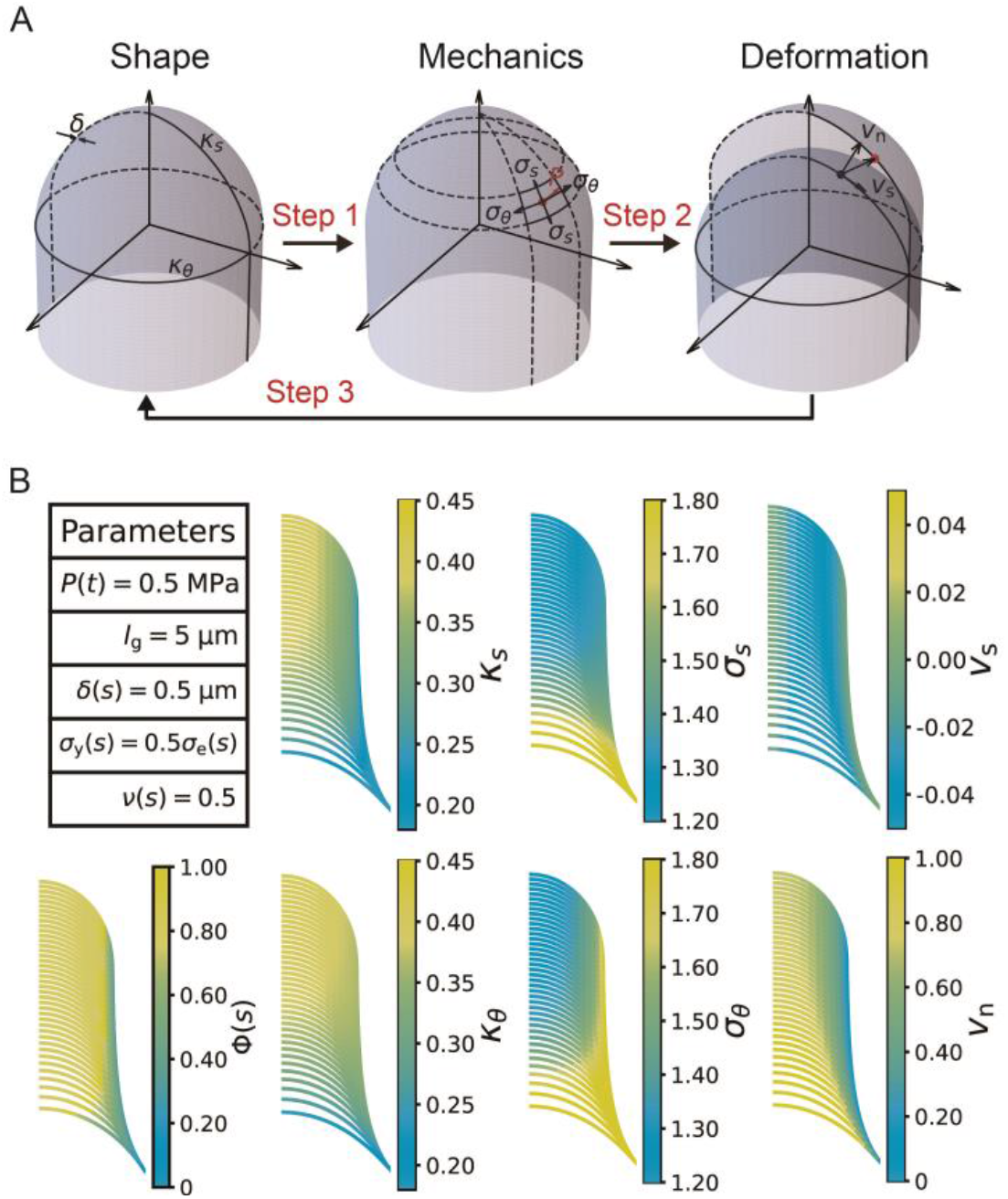
Simultaneous evaluation of the shape, mechanics, and deformation of tip-growing cells. (A) Schematic representation of the model. The mechanics variable *σ* is determined from the shape data (*κ, δ*), through mechanical equilibrium with turgor pressure *P*. The deformation variable 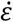 is determined from the mechanics input. (B) Evaluated shape, mechanics, and deformation variables and wall extensibility.

As the model includes the shape, mechanics, and deformation of the cell at each time step, it is reasonable to describe all of these simultaneously, as shown in Fig. 2B. Using the parameters in Fig. 2B, we obtained the shape information (*κ*_s_, *κ*_θ_), mechanical information (*σ*_s_, *σ*_θ_), and deformational information (*v*_n_, *v*_t_) for a typical tip-growing cell with radius 1 (a.u.). This simultaneous evaluation is important because it incorporates the mechanical and deformational events simultaneously with the corresponding shape change.

### Cosine-type wall extensibility results in the formation of a hemisphere-like tip shape

To further investigate the characteristic features of the model, we investigated the strain profile derived from stress input, turgor pressure, and cell wall extensibility. The strain profile is defined as the curvilinear coordinate system *S*, where *S* = 0 at the tip and *S* = *s* at position *s* (Fig. 3A). The cell wall extensibility Φ(*S*) is the degree of surface growth, as presented in Fig. 3D, where the magnitude at *S* = 0 is denoted by Φ_0_ and the range of the extension zone is denoted by *l*_g_. Based on this strain profile, we explored the spatial distribution of cell wall extensibility. As the zygote maintains an approximately hemispherical shape (Fig. 3A–C and Figs. S1−S3), the distribution of surface growth should reflect this shape change (the current hemisphere to the next hemisphere).

**Fig. 3.**
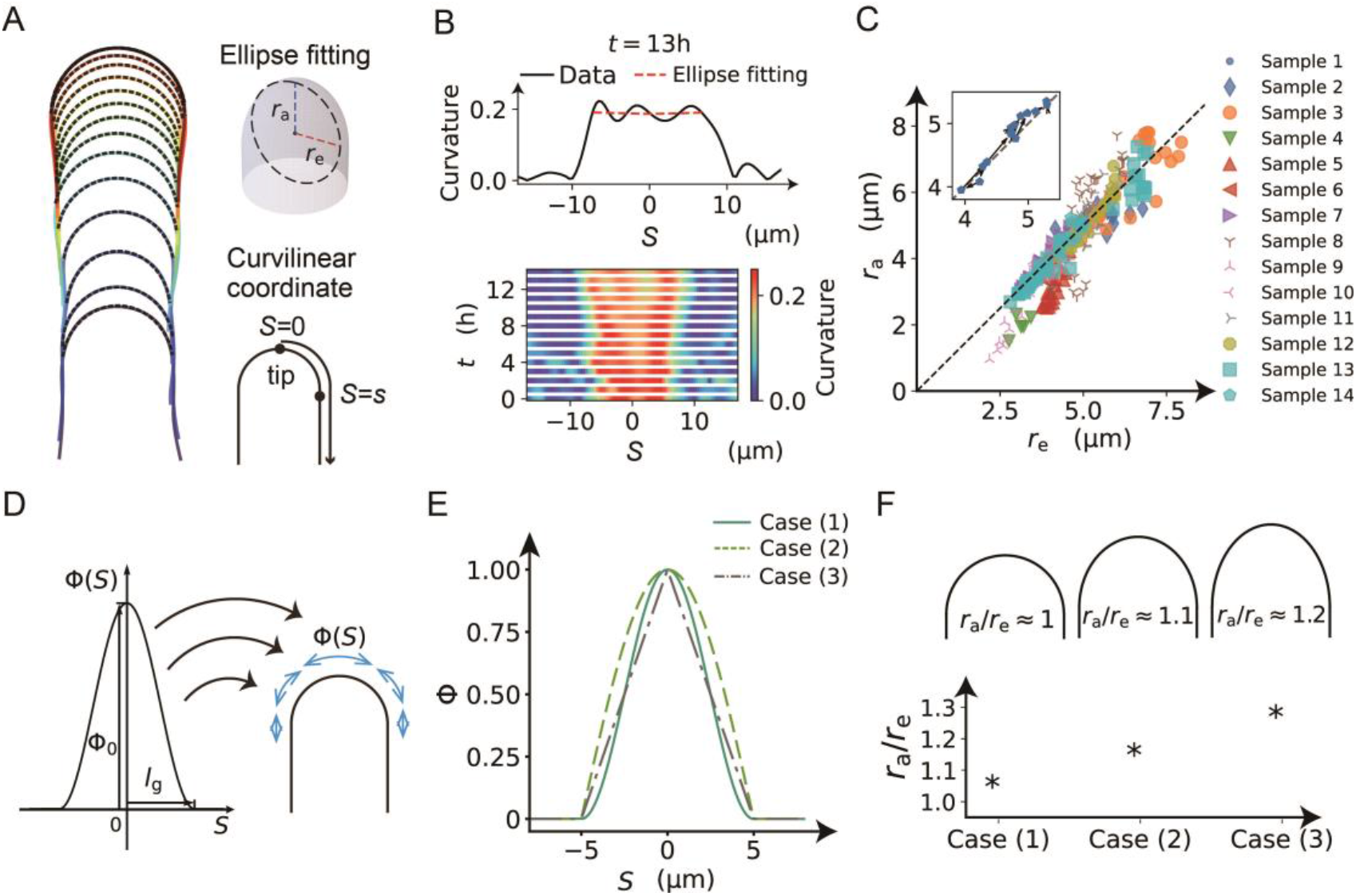
Cosine-type wall extensibility model showing a hemisphere-like shape change during cell elongation. (A) Left panel shows cell contours of a zygote with temporal color code from blue to red with ellipse fitting (black dashed lines). Right panels show schematic illustrations of ellipse fitting and curvilinear coordinate with *S* = 0 at the tip. (B) Top panel shows the curvature profile as a function of S and that from ellipse fitting (red dashed line). Bottom panel shows the spatio-temporal kymograph of the curvature. (C) Plot of *r*_e_ and *r*_a_ with the diagonal *r*_e_ = *r*_a_ (dotted line) based on the dataset in Kang *et al*., 2023. (D) Spatial distribution of cell wall extensibility. Greater magnitude of cell wall extensibility indicates a high strain rate at the extension zone. (E) Three different profiles of cell wall extensibility were considered: case (1) Φ(*S*) = cos(π*S*/2*l*_g_), case (2) Φ(*S*) = cos^2^(π*S*/2*l*_g_), and case (3) Φ(*S*) = 1 − *S*/*l*_g_. (F) Ellipse fitting revealed that the aspect ratio of tip shape becomes close to 1.00 in the case of a cosine-type profile. The values for the three examples are *r*_a_/*r*_e_ ≈ 1.01 for cosine, *r*_a_/*r*_e_ ≈ 1.11 for square of cosine, and *r*_a_/*r*_e_ ≈ 1.23 for linear function.

The time-averaged velocities of the three example zygotes reported by Kang *et al*. (2023) are presented in supplementary Fig. S4, where the tip has a maximum growth rate that gradually decreases as the meridional distance increases. We also confirmed that the curvature away from the tip did not change significantly over time for all the samples shown in supplementary Fig. S5. This indicates that the zygote cells grow in the manner of tip growth with a hemisphere-like shape. Acknowledging that early studies (Green & King 1966; Green 1969; Dumais *et al*., 2004) demonstrated that the hemispherical shape is sufficient for cosine-type wall extensibility in our model, the hemispherical shape is ensured by the cosine-type wall extensibility only if the growth is transversely isotropic (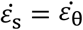, see Methods). Therefore, we sought different types of Φ(*S*) without assuming transversely isotropic growth. We employed three different formulations: case (1) Φ(*S*) = cos(π*S*/2*l*_g_), case (2) Φ(*S*) = cos^2^(π*S*/2*l*_g_), and case (3) Φ(*S*) = 1 − *S*/*l*_g_, as shown in Fig. 3E. We then quantified the ellipse-fitting parameters (*r*_a_, *r*_e_) using the radial half axis *r*_a_ and the circumferential half axis *r*_e_ for our model (Fig. 3F). By definition, the aspect ratio *r*_a_/*r*_e_ > 1 corresponds to a tapered shape, *r*_a_/*r*_e_ = 1 corresponds to a hemispherical shape, and *r*_a_/*r*_e_ < 1 corresponds to a flattened shape. As shown in Fig. 3D, the aspect ratios obtained were close to the value *r*_a_/*r*_e_ ≈ 1.01 for (1), *r*_a_/*r*_e_ ≈ 1.11 for (2), and *r*_a_/*r*_e_ ≈ 1.23 for (3). The actual data for the aspect ratio of cell shape were approximately equal to 1.00, corresponding to a hemisphere-like shape, indicating that the cosine-type wall extensibility profile is well fitted to tip-growing cells in plant zygotes.

### The hemisphere-like cell growth model exhibits a normal growth direction around the cell tip

To clarify what happens to the growth direction in the model with a hemisphere-like shape, we investigated the directional angle *ψ*, which measures the deviated angle from the normal surface direction to the direction of surface point velocity (Fig. 4A). The color diagrams of *ψ* as a function of *S* show that *ψ* is approximately equal to 0 regardless of *S*, which indicates a normal growth direction around the cell tip (Fig. 4B). By contrast, the angle *ψ* for cases (2) and (3) shows positive values in the subapical regions (Fig. 4C and Fig. 4D), indicating that the growth direction is not normal. Therefore, the resulting shape in cases (2) and (3) becomes more tapered. Therefore, we concluded that the cosine-type wall extensibility leads to the normal growth direction.

**Fig. 4.**
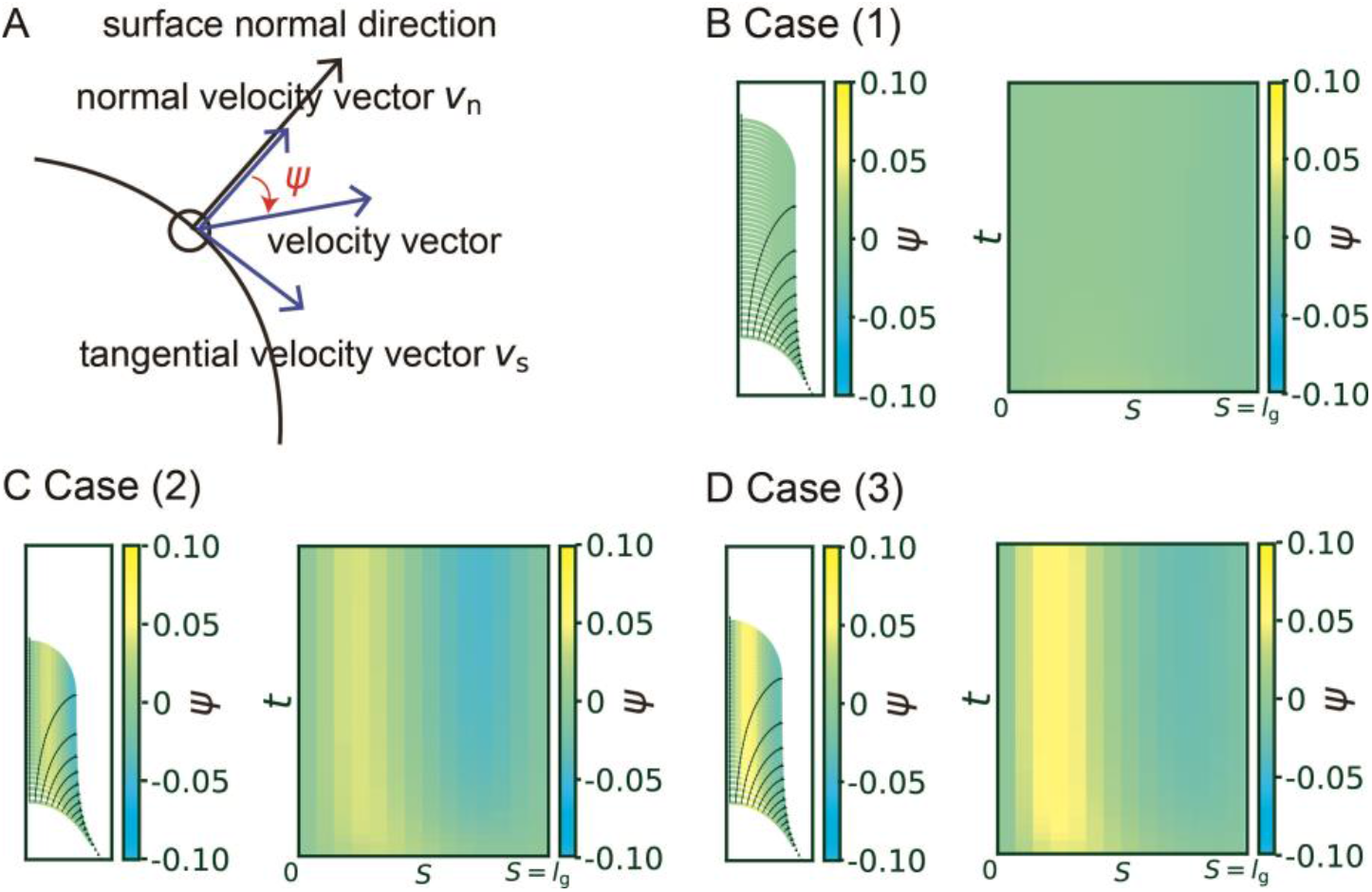
Hemispherical shape results from the normal growth direction during cell elongation. (A) Definition of the growth angle *ψ*. (B–D) Growth trajectories of selected points (black lines) with color code *ψ* are shown in the left panel, and spatio-temporal plots of the corresponding color code *ψ* are shown in the right panel for case (1) Φ(*S*) = cos(π*S*/2*l*_g_) (B), case (2) Φ(*S*) = cos^2^(π*S*/2*l*_g_) (C), and case (3) Φ(*S*) = 1 − *S*/*l*_g_ (D).

### An elongating zygote can be reconstructed computationally from cell contour data alone

As described above, we obtained a mathematically supported logical connection between cosine-type wall extensibility and the normal growth direction for hemisphere-like zygote shapes. The relationship points to the possibility of inferring model parameters from only cell contour data using such a constraint for wall extensibility. In our previously reported live-imaging time sequence of zygote plasma membrane markers, we obtained cell contours using the cell contour–based coordinate normalization (CCN) method (Kang *et al*., 2023). Using the cell contour, we quantified the so-called morphospace 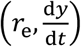, where d*y*/d*t* is the growth rate in the y-axis (Fig. 5A). Among the previously reported samples (Kang *et al*., 2023), parameters were distributed in the range *r*_e_ ∈ [3,5] and d*y*/d*t* ∈ [1,5]. For the sake of simplicity, we considered the time-averaged values for each sample (Fig. 5B). Based on perturbation analysis of *P* and *l*_g_ in the simulations (Fig. 5C), we noticed that *P* only affects the growth rate, while *l*_g_ predominantly changes the radius. Therefore, we classified all the samples into group 1 or group 2 and fitted the parameter set (*P, l*_g_) for the sample-averaged values of (*r*_e_, d*y*/d*t*) denoted by Examples 1 and 2, respectively. To further confirm what happens for intermediate parameters, we included Example 3, with almost the same growth rate as Example 1 and almost the same radius as Example 2. Using typical values for each quantity (*r*_e_, d*y*/d*t*) with the previously estimated range of *P* (0.3– 1.0 MPa) (Cosgrove, 1993; Lintilhac *et al*., 2000; Radotić *et al*., 2012) (Fig. 5C), we applied an exhaustive search of *l*_g_ parameters that matched the data and obtained the fitted parameter *l*_g_. With these parameters, we reconstructed the elongating zygote computationally (Fig. 5D). Note that we assumed a constant cell wall thickness (*δ* = 0.5 μm) in this study based on the acknowledgement that the parameter Φ_0_ was affected by *δ*.

**Fig. 5.**
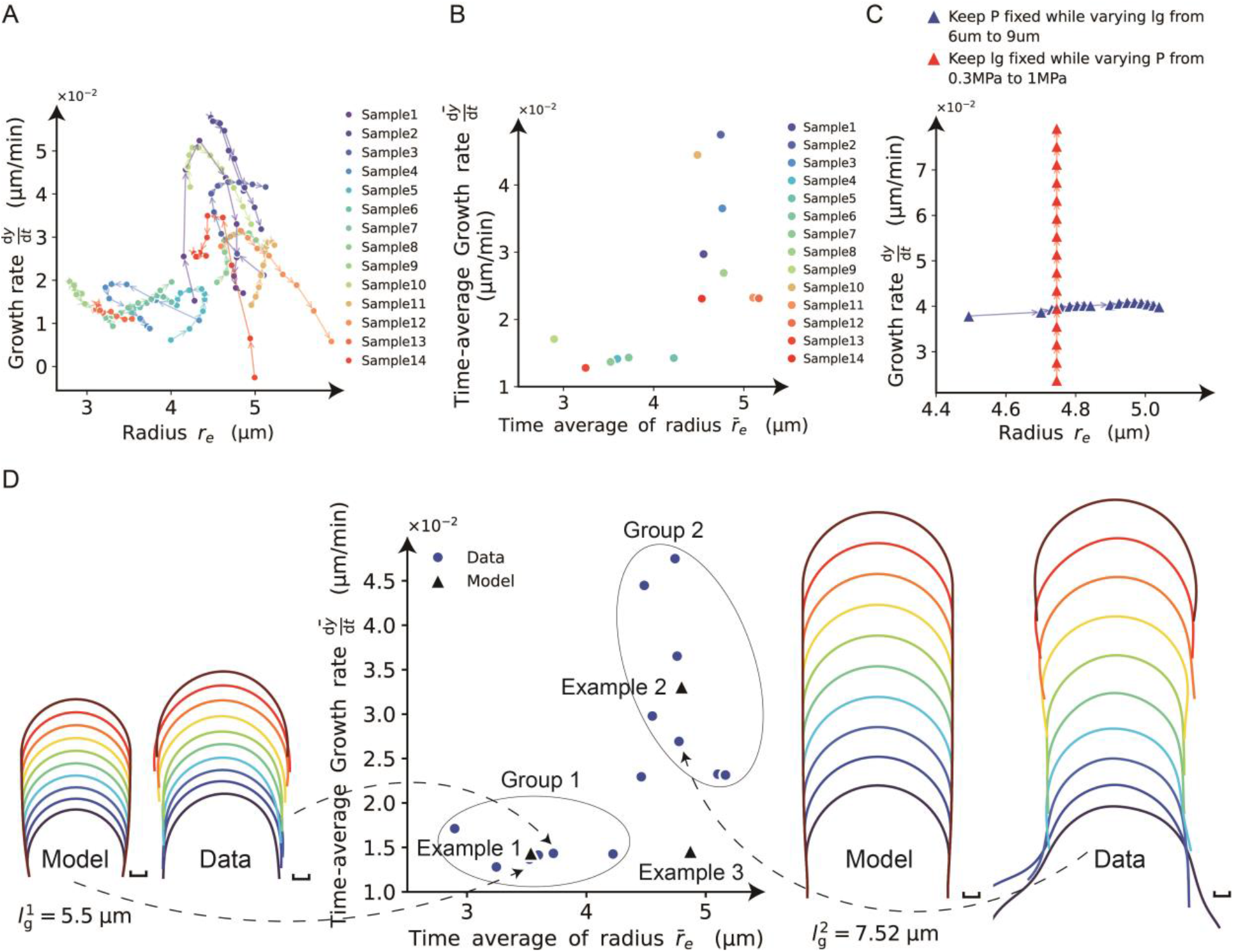
Reconstruction of model parameters using only cell contour data. (A) Morphospace analysis using the circumferential half axis *r*_e_ and the growth rate in the y-axis d*y*/d*t*. (B) The time-average of the growth rate and the radius for each sample. (C) Perturbation analysis of *P* and *l*_g_ in the mechanical simulations. (D) The samples are classified into group 1 or group 2, with the sample-averaged values of (*r*_e_, d*y*/d*t*) denoted by Examples 1 and 2, respectively. The left panel shows the reconstructed model for Example 1 and one data point from the contour data for group 1. The right panel shows the reconstructed model for Example 2 and one data point from the contour data for group 2.

To summarize, we were able to reconstruct the elongating zygote computationally from only the cell contour data, with the parameter ranges inherited from the data range.

### Non-dimensional control parameters (*α, β*) characterize tip-growing behavior

Through the above model reconstruction, we determined effective parameters as follows. First, as the growth rate is affected by *P*, Φ_0_, and *l*_g_, we defined the cell growth sensitivity as

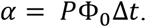

This is a non-dimensional parameter that reveals the vertical growth length (Fig. 6A). As the parameter *r*_e_ is another characteristic of the zygote cell, we can also define the constant

**Fig. 6.**
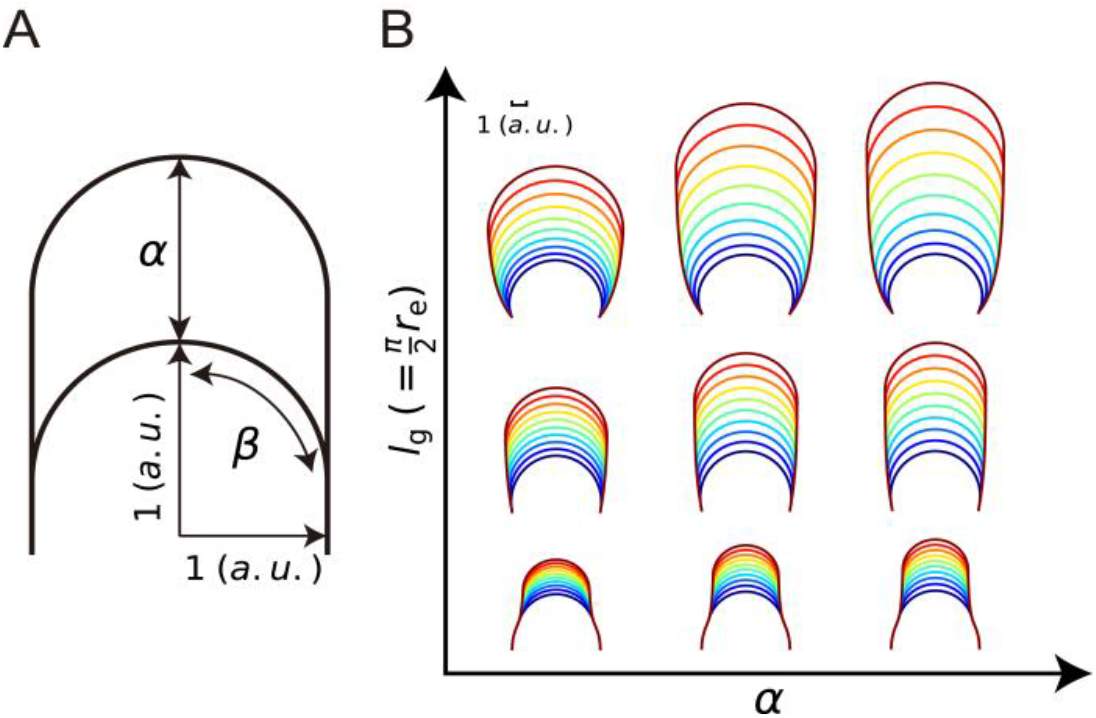
Tip-growing behavior is controlled by non-dimensional parameters (*α, β*). (A) Schematic illustration of the non-dimensional parameters *α* and *β*. (B) Morphospace for the elongating zygote where the stable elongating cell shape depends only on the non-dimensional parameters (*α, β*).

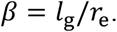

This is the other non-dimensional parameter and is the ratio between meridional growth length and hemispherical half axis length (Fig. 6A). Under the hypothetical condition of the cosine-type wall extensibility profile, *β* = π/2. By definition, the parameters *α* and *β* are independent of cell size; therefore, these can be used to sufficiently characterize tip-growing behavior, as shown in Fig. 6B.

## Discussion

The viscoelastic-plastic deformation model developed in this study explores three new aspects of cell elongation analysis. (1) Simultaneous evaluation of shape, mechanics, and deformation, which will be a powerful tool for understanding multiple factors during cell elongation. (2) Quantitative verification of normal growth direction to produce a hemisphere-like shape. (3) Model reconstruction using only the cell contours derived from live-imaging data. Some possible future directions are summarized below.

In the field of plant mechanobiology, mechanical measurement is critical, such as in atomic force microscopy (Beauzamy *et al*., 2015; Tsugawa *et al*., 2022) and in mechanical inference from stem shape (Nakata *et al*., 2018). However, these methods are only applicable to the surface cells and external shapes of organs, whereas the mechanics of developmentally important cells, such as the zygote cells within the seed and lateral root primordia or vascular cells inside the root, have not yet been fully examined. Using our data–model combined method, we defined the model parameters using only the quantification of cell contours. Therefore, our method can precisely infer cellular mechanics, which cannot be determined solely by current mechanical measurement techniques, opening a new avenue for studying cellular mechanics not only in plant developmental biology but also in other multicellular systems including animal cells.

Considering the Lockhart equations (Lockhart, 1965a, 1965b), which guide reconstruction of the plastic deformation of plant cells, the cosine-type distribution Φ(*S*) and its effect on morphology can be thought of as a spatial example of plastic deformation relating to hemisphere-like tip shape. Biological events regulating cosine-type distribution should include the heterogeneous distribution of microtubule ring structure (Kimata *et al*., 2016), where the microtubule-associated cell wall deposition acts as a mechanical hoop to regulate hemisphere-like tip shape. Furthermore, since our model includes viscoelastic deformation, it also considers the later-proposed viscoelastic-plastic deformation (Ortega, 1985). As revisited by Green *et al*. (1971), a possible first approach for reconstructing plastic deformation including viscoelasticity is to use models that take into account a threshold level of viscoelasticity. The present model precisely reflects the understanding that cell wall loosening occurs before water uptake (Cosgrove, 1985, 1987), implying that the parameter Φ(*S*) is the most important wall parameter involved in morphology, stress, and strain. In addition, we identified the non-dimensional indices *α* and *β* that fully characterize the dynamics of our tip-growing cells.

In summary, we developed a data-driven reconstruction method of mechanical model for studying cell elongation. This method represents a promising tool for studying the growth mechanisms and biological connections between morphology, mechanics, and deformation, paving the way for understanding tip-growing cells.

## Supporting information

QPB_Kang_sup_ver7.pdf

## Acknowledgements

The authors thank Koichi Fujimoto and his lab members (Hiroshima University) and Takumi Higaki and his lab members (Kumamoto University) and Yusuke Kimata (Tohoku University) for helpful discussions.

## Author Contributions

Z.K. and S.T. conceived and designed the study. Z.K. and S.T. developed and implemented models and algorithms. H.M., S.N. and M.U. carried out data acquisition from live cell imaging. Z.K. and S.T. wrote the manuscript. Z.K., T.N., S.T., M.U. and Y.I. edited and reviewed the manuscript. S.T. and M.U. provided research funding. All authors have read and approved the manuscript.

## Data Availability Statement

The data supporting the findings of this study are available from the author, Zichen Kang, upon a reasonable request. The code used in this study are available on GitHub (https://github.com/blues0910/A-viscoelastic-plastic-deformation-model-of-hemisphere-like-tip-growth).

