## Supplementary material for "A viscoelastic-plastic deformation model of hemisphere-like tip growth in Arabidopsis zygotes": QPB_Kang_sup_ver7.pdf

### Supplementary Figures S1-S5

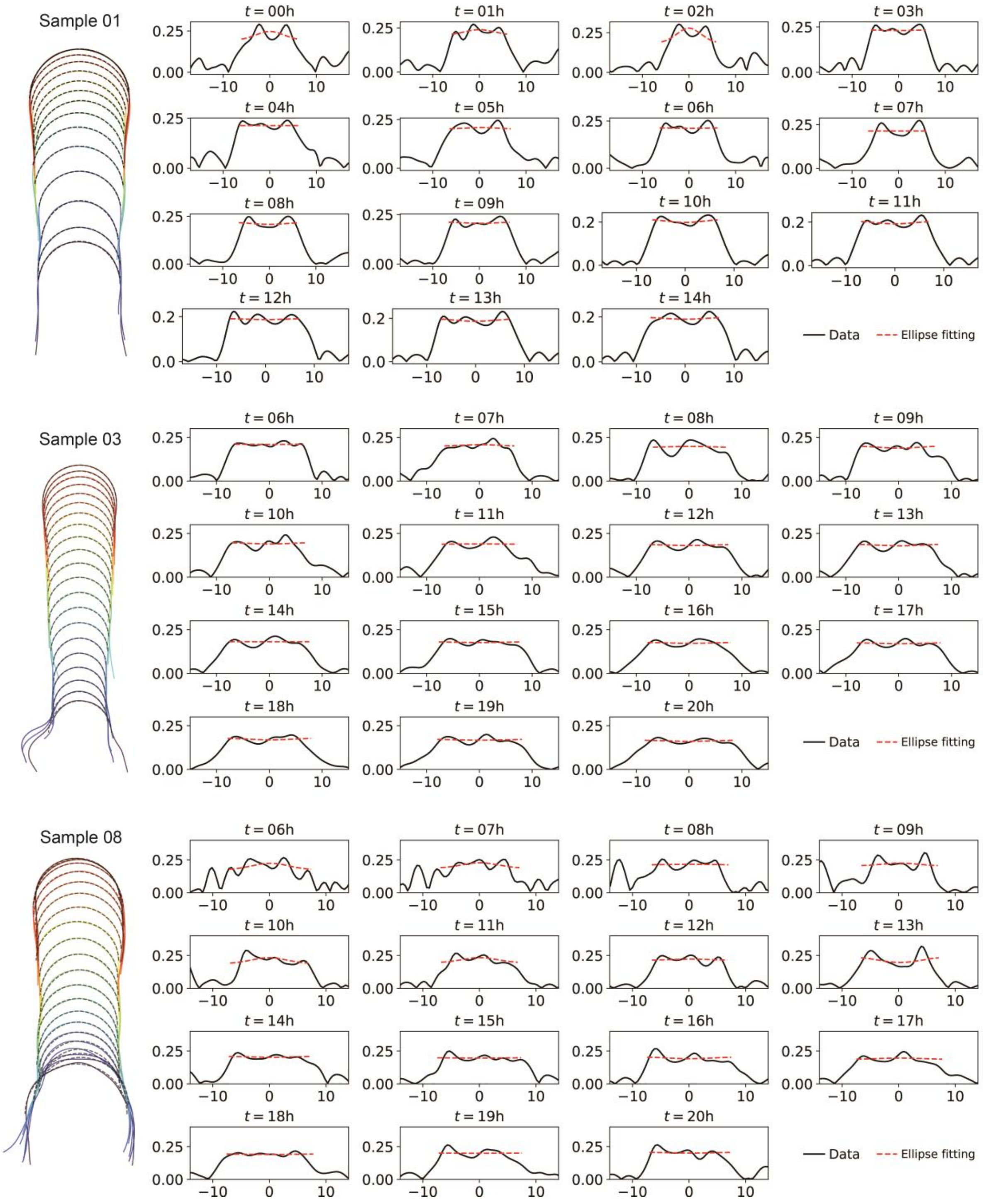

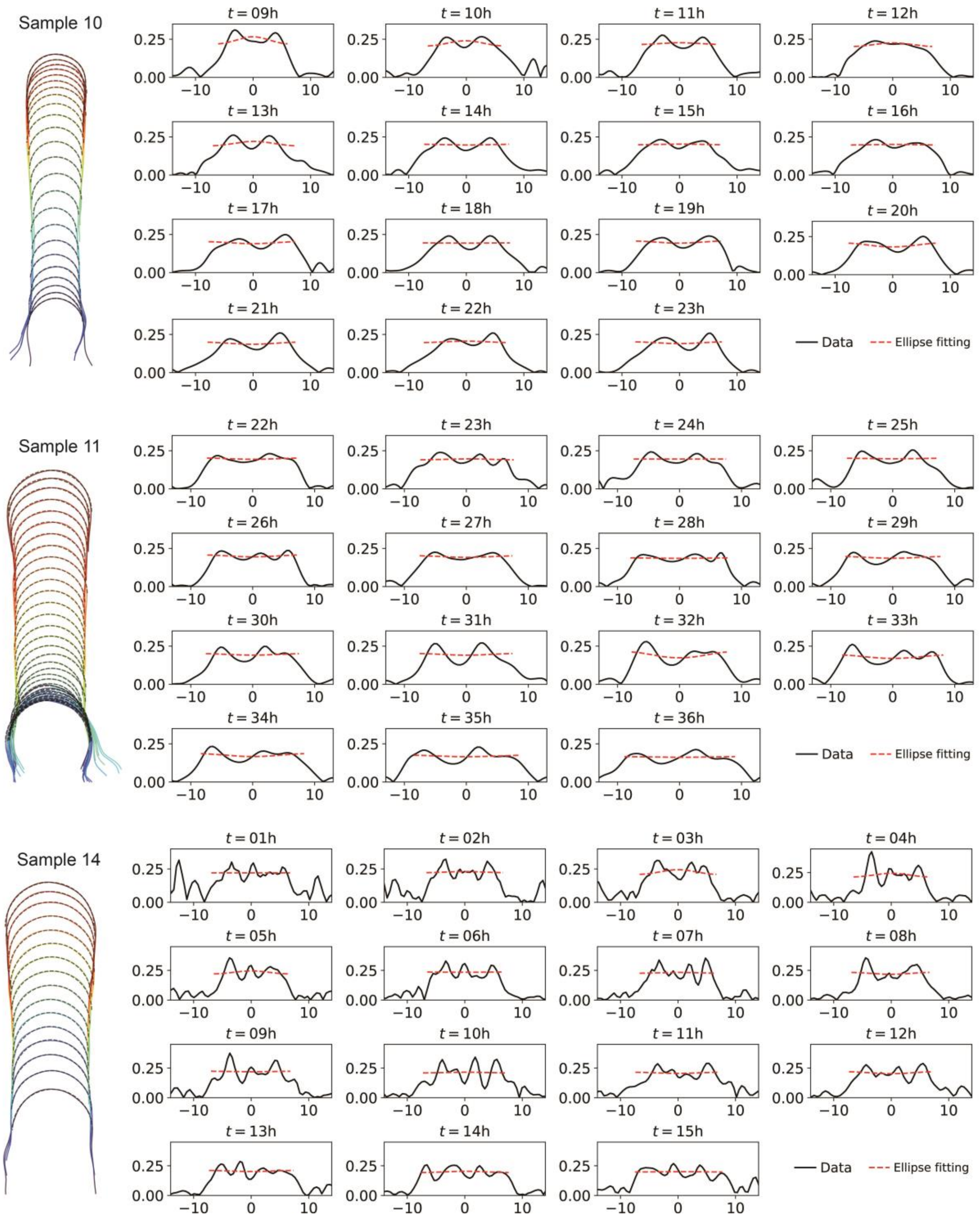


**Supplementary Fig. S1.** Verification of hemisphere-like shapes of elongation zygote tips using ellipse fitting for the dataset in Kang *et al*., 2023. The left panels show zygote cell contours during elongation with temporal color code from blue to red. The black dashed lines represent the upper halves of the ellipses fitted to the tip contours. The right panels show the curvatures of the tip contours for every time point with fitted results (red dashed lines).

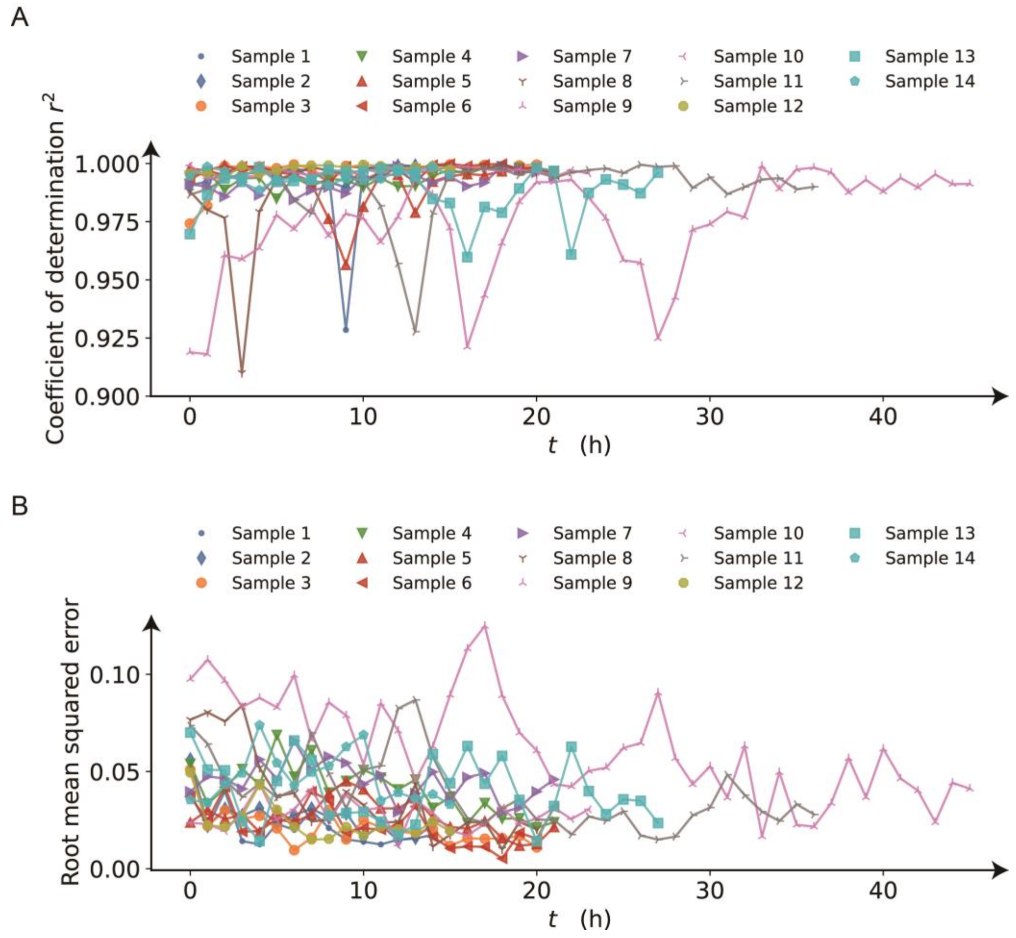


**Supplementary Fig. S2.** Evaluation of fitting errors of the ellipse to the cell contours in supplementary figure S1. (A) Coefficient of determination of the fitted ellipse for each sample as a function of time. (B) Root mean squared error between the curvatures of the fitted ellipses and the curvatures in data.

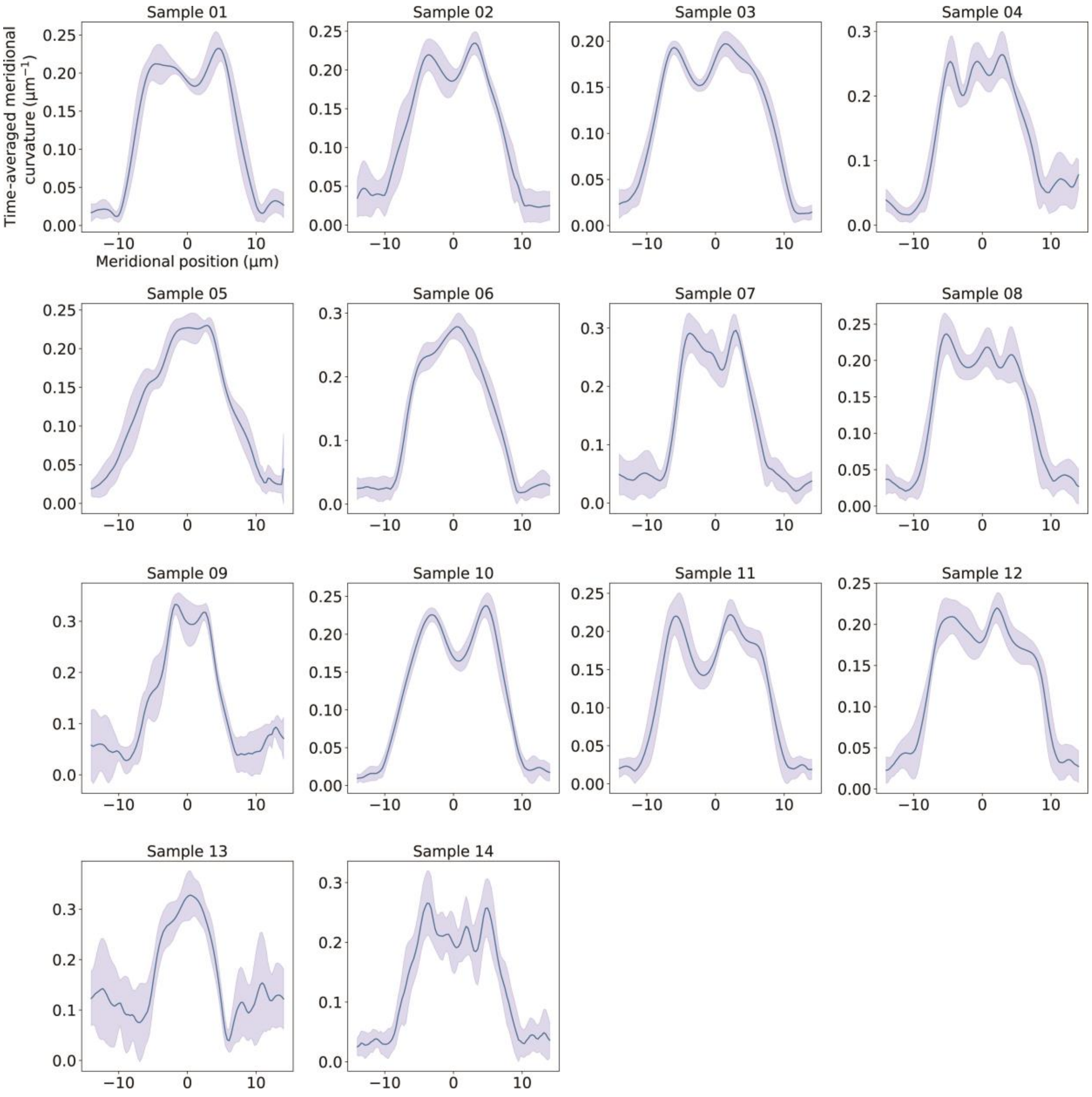


**Supplementary Fig. S3.** Time-averaged meridional curvature of zygote tip as a function of meridional coordinate for the dataset in Kang *et al*., 2023. The blue line and shed regions around the lines stand for the time average and the standard deviations.

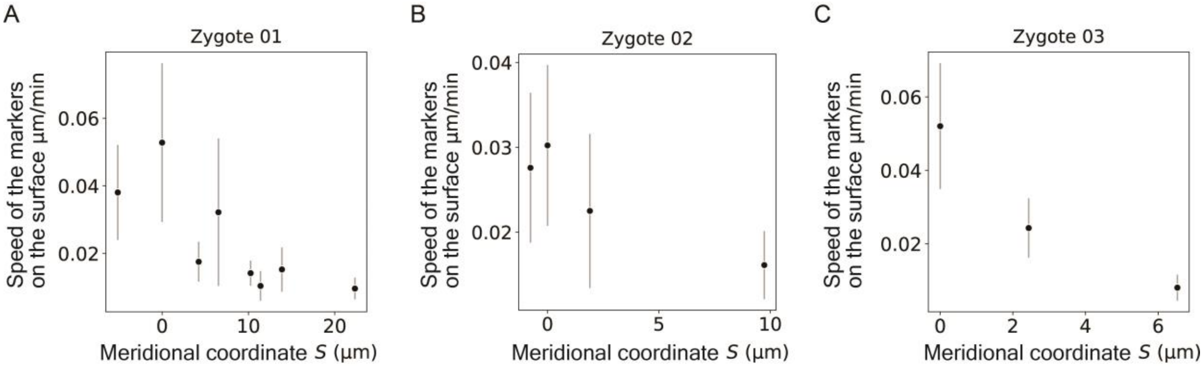


**Supplementary Fig. S4.** Time averaged speed of the markers on the cell surface based on the dataset in Kang *et al.*, 2023. The black points and bars stand for the time average and the standard deviations.

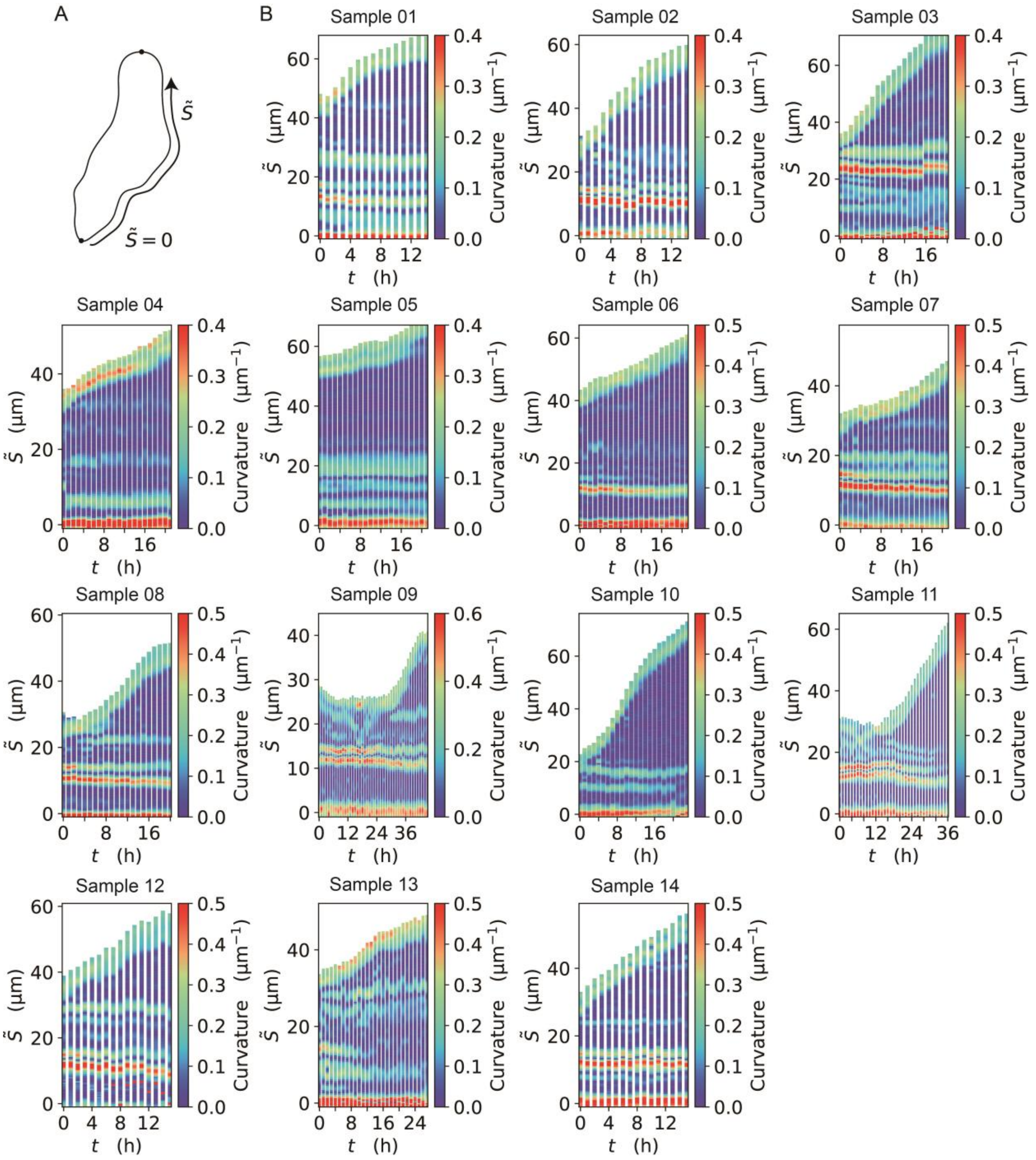


**Supplementary Fig. S5.** Quantitative evidence that the zygote dominantly grows around tip region over time. (A) Schematic illustration of the curvilinear coordinate $\tilde{S}$ where $\tilde{S} = 0$ at the bottom. (B) Spatio-temporal kymograph of curvature based on the dataset in Kang *et al.*, 2023, showing that the curvatures of the regions away from the tip remain constant over time.
